# Reinforced Random Walker meets Spike Timing Dependent Plasticity

**DOI:** 10.1101/168401

**Authors:** Mohammadreza Soltanipour, Hamed Seyed-allaei

## Abstract

We blended Reinforced Random Walker (RRW) and Spike Timing Dependent Plasticity (STDP) as a minimalistic model to study plasticity of neural network. The model includes walkers which randomly wander on a weighted network. A walker selects a link with a probability proportional to its weight. If the other side of the link is empty, the move succeeds and link’s weight is strengthened (Long Term Potentiation). If the other side is occupied, then the move fails and the weight of the link is weakened (Long Term Depression). Depending on the number of walkers, we observed two phases: ordered (a few strong loops) and disordered (all links are alike). We detected a phase transition from disorder to order depending on the number of walkers. At the transition point, where there was a balance between potentiation and depression, the system became scale-free and histogram of weights was a power law. This work demonstrate how dynamic of a complex adaptive system can lead to critical behavior in its structure via a STDP-like rule.

## 1. Introduction

Learning is among the most underlying traits of the brain, which is carried out through strengthening and weakening of synapses between various neurons (synaptic plasticity) [1, 2]. The case of a neuronal activity resulting in the strengthening of a specific synapse is called Long Term Potentiation (LTP), and in the case of its weakening, Long Term Depression (LTD) [3].

Spike Timing Dependent Plasticity (STDP) is an asymmetrical form of Hebbian learning that models synaptic plasticity based on the time difference between firing of pre and post-synaptic neurons [4, 5, 6]. In the most common form of the STDP model, for which there exist many experimental confirmations, pre-synaptic neuron spiking prior to a post-synaptic neuron results in LTP, and spiking of a pre-synaptic neuron subsequent to the spiking of a post-synaptic neuron results in LTD [7, 8, 9].

A lot has been done to study the effects of STDP on the evolution and on the properties of the final structure of neural networks [10, 11, 12, 13, 14]. In this paper, by proposing a model of reinforced random walkers, we tried to reconstruct the main properties of STDP.

Reinforced Random Walkers (RRW) are random walkers on a network, such that their passage through a link increases the chance that they will pass through the same link in the future [15]. RRW have been used to model the behavior of cells whose passage through a specific point causes the chemical environment to adjust to the transmission of the rest of the cells [16]. To illustrate, this is used to model the movement of myxobacteria (a bacteria that lives in soil) [17] and the migration of endothelial cells during tumor-induced angiogenesis [18, 19, 20].

In the original implementation, the walkers only reinforce the links as they pass but in a recent model, Mehraban and Ejtehadi introduced an RRW with two kinds of walkers, the first reinforces the connections and the second plays a weakening role. Depending on the ratio of walkers, they observed three phases including, ordered, disordered and transition phase [21].

In the following sections we will show that how our model, although much simplified, possesses the main characteristics of STDP and report how a network subject to this model evolves and turns into its final structure.

Many evidences suggest that the brain operates on a critical point or somewhere close to it, by observing scale-free behavior in variables such as avalanche sizes [22, 23, 24], a benefit of which is the maximization of information processing [25].

It has been suggested that the Synaptic Plasticity Plays a crucial role in getting to the critical point [26, 27, 28]. Most of these studies address functional criticality, when the activity of the network shows the sign of critical behavior [29, 30].

In the current research, we are focused on the structure of the network and we have shown that at a specific point when, what we interpret as LTP and LTD play an equal role in the evolution of the system, the system finds itself in the critical point in respect to the synaptic weights. This point guides us towards a new idea regarding how the brain approaches the critical point.

## 2. Model

Our model is a reinforced random walk on directed network without any self loops. We start by a fully connected network, for which all the weights are equal. We then insert some walkers into the network. Each node has at most one walker (walkers are fermions), which until picked, seats on the node. Each network has three parameters: number of its nodes (*n*), number of its walkers (*m*), and evolution rate constant (*a*), where *a* = 1 + *ϵ* and *ϵ* ≪ 1. At each time step, we randomly pick a node and if it contains a walker, we select one of its neighbors with a probability of *p_ij_*, where *P* is the weight matrix of the network. After selecting the walker’s destination, two cases may happen when:

i. Selected destination node *j* is not occupied by any other walkers, where the motion takes place and the selected link will be strengthened:

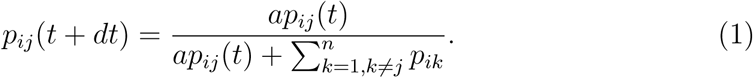
ii. Selected destination node *j* is already occupied by another walker, where the motion dose not take place and the selected link will be weakened:

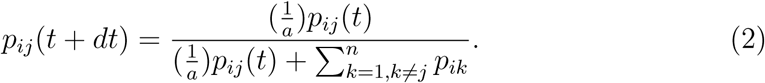

In our simulations the evolution rate is assumed to be a fixed value of *a* = 1.1 (*ϵ* = 0.1).

The weight of outgoing links from a given node sums to one, so, in addition to strengthening (weakening) of the link’s transition probability, the transition probabilities of the remaining links associated with the departure node is weakened (strengthened).

We observed the evolution of the network over time. The initial form of the network was a fully connected network with equal weights. However, due to the self-organization property of the model, final properties of the network after reaching the steady state are quite independent of its initial conditions – the initial values of the transition probability matrix *P* and the initial place of walkers.

We assume that the system is in the steady state when the average Shannon entropy per node has reached a constant value and will remain unchanged thereafter. The entropy per node is defined by [31]:

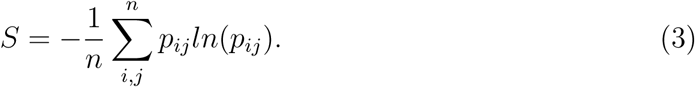

In our model, each node plays the role of a neuron and the existence of a walker in a node translates to activity *(spiking)* of that neuron. The model in question, albeit very simple, possesses the main properties of the strengthening and weakening of a synapse:

i. LTP and LTD are reconstructed by successful and unsuccessful movement attempts: Each time a walker succeeds (post-synaptic neuron is stimulated by the pre-synaptic neuron and spikes), the synapse between two neurons (the link between the two nodes) is strengthened (LTP). And each time a walker fails (pre-synaptic neuron tries to stimulate the post-synaptic neuron just when the latter is active), the synapse between two neurons is weakened (LTD).
ii. Most of the time Hebbian learning requires synapses competition [32, 11]. In our model, as we saw, not only a successful move increases the transition probability of that link, but it also decreases the transition probability of other links associated with the departure node.

## 3. Results

We simulated different networks with different number of nodes *n* and walkers *m*, and judging by the results, the final sate of the network was merely a function of walker density 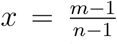 (to exclude the current selected walker and node we subtracted *m* and *n* by one). We used entropy per node (Eq.3) as an order parameter. Sweeping *x* from zero to one and monitoring the final entropy of the system, we observe a transition phase which takes place at exactly 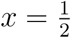 (Fig.1).

**Figure 1.**
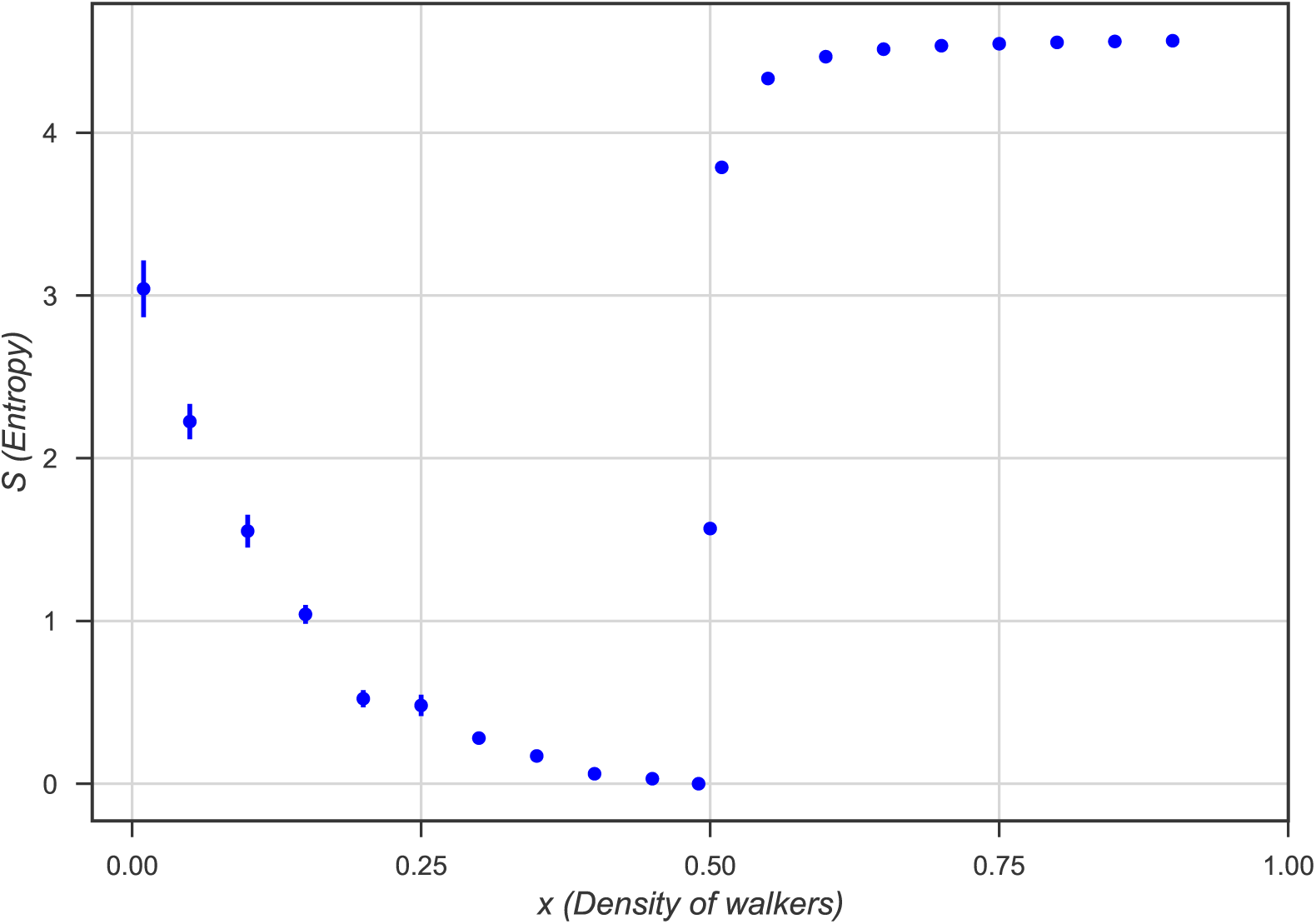
Entropy of the system at steady state for network of size 101, with respect to density of the walkers, 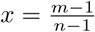. The phase transition occurs exactly at 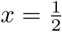.

Depending on the value of *x*, the system falls to one of the three distinct phases: ordered, disordered and transition phase.

### Ordered phase

Depending on the value of *x*, the system falls to one of the three distinct phases: ordered, disordered and transition phase.

### Ordered phase

For *x* < 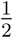, the system reaches the ordered phase. In this phase, a few loops form in the network and all of the walkers move throughout them. The emergence of loops indicates that the transition probability of the links for each node *i* is as follows: There is a node *k* which *p_ik_* → 1, and for all others (*i* ≠ *k*), *p_ij_* → 0. A sample of this phase can be seen in Fig. 3. If every node of the network is incorporated in a loop, the entropy of the system will be equal to zero (Eq.3). But as it can be seen in Fig.3, for *x* < 0.4 we have an entropy greater than zero, which increases upon decreasing *x*. The reason for the emergence of such behavior is that some nodes would not evolve as a result of decreasing *x*, and in practice they do not take part in any loops. Indeed, after a while, the system creates a few loops which attracts all of the walkers and leave no walker for other nodes to evolve. This behavior is limited upon the increase of *x*, for as the number of walkers increase, they are going to need loops which incorporate a greater number of nodes for them to move through, because, if there are not many nodes participating, the number of the failed movement begins to increase, which effectively ruins the loop. The aforementioned behavior can be observed according to the histogram of transition probability (Fig.2). As is evident, with the walkers increasing, we have more links with the probability 1 and less links with probabilities between 0 and 1.

**Figure 2.**
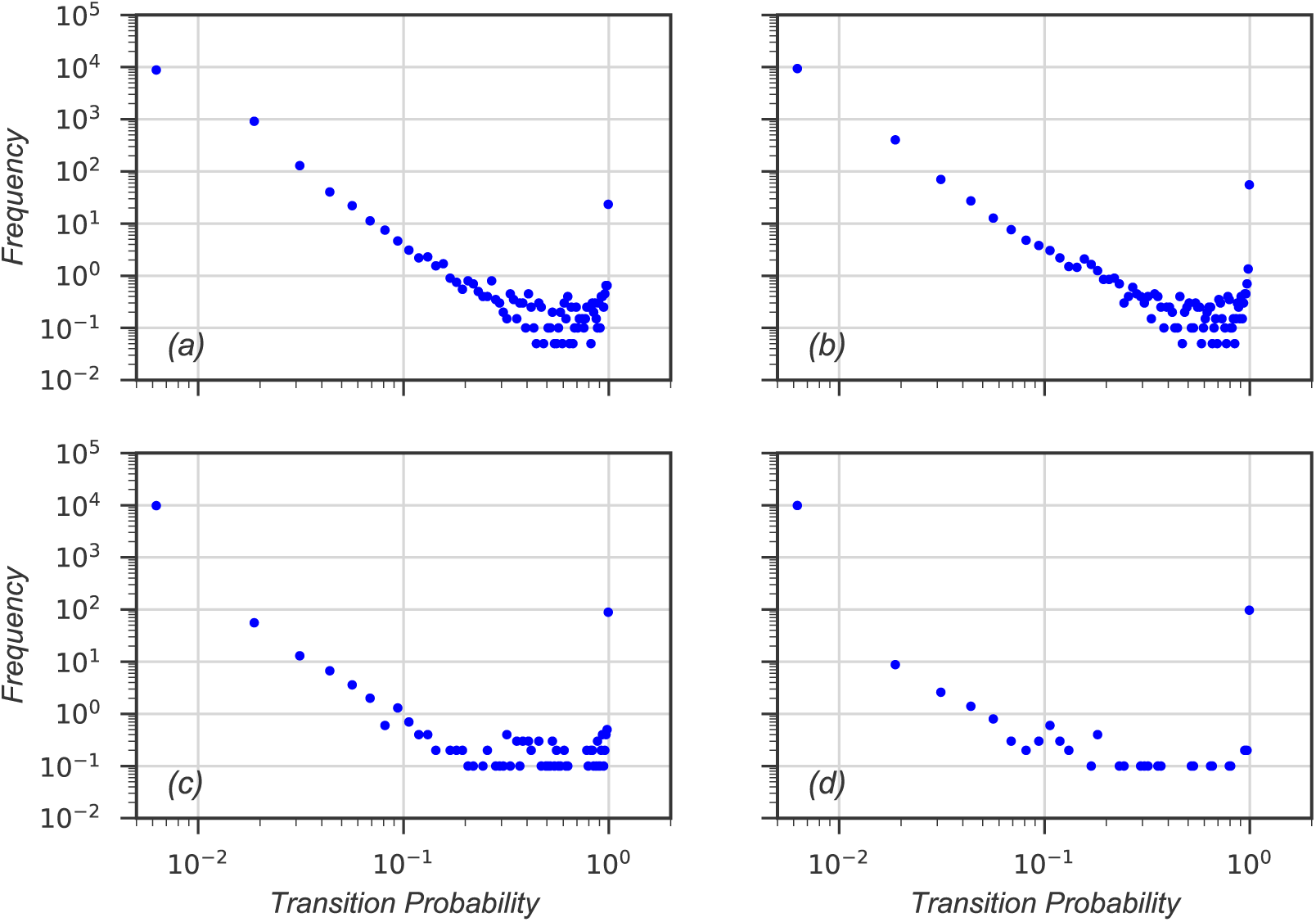
The histogram of transition probability for four different densities in the ordered phase in the network of *n* = 101 nodes. a)*x* = 0.01, b)*x* = 0.10, c)*x* = 0.30 and d)*x* = 0.40. The number of the nodes participating in the loops grows as x increases, and as a result there exists more nodes that have 1 for the transition probability of one of their links, and 0 for the rest.

**Figure 3.**
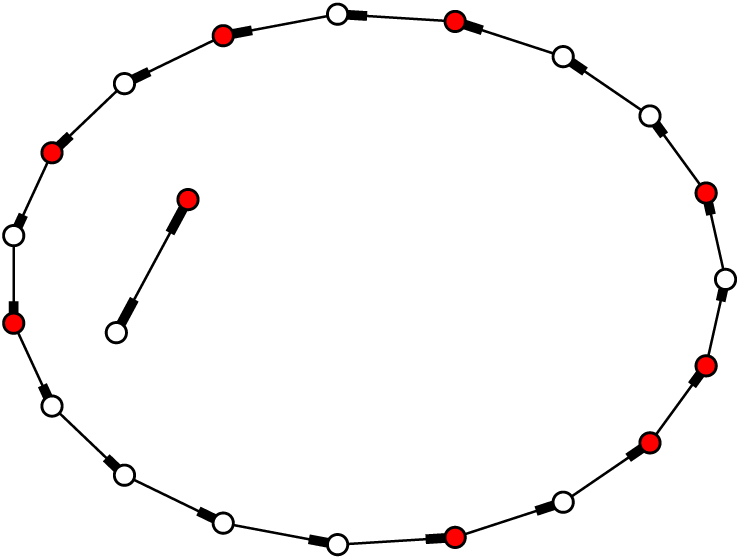
The steady state structure of a sample network with 21 nodes and 9 walkers (ordered phase). All the links have a weight close to 1. The formation of loops are evident.

### Disordered phase

For *x* > 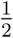, the system will reach the disordered phase, such that, as a result of more than half of the nodes being occupied, at each time step the probability of weakening is greater than the probability of strengthening of the selected link, and the decrease in the transition probability of a specific link results in the same state, similar to that of other links. Eventually, the elements of the probability ratio matrix reduce to a symmetrical form, which translates to equal transition probability for all links; Therefore, we expect that in this phase, all the links emerging from a node have equal probability, 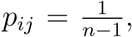 for *i* ≠ *j*. In this case the entropy reaches its maximum (Eq.3), and as we can see in Fig.1 and Fig.4, the results agree with what we have expected from the model.

**Figure 4.**
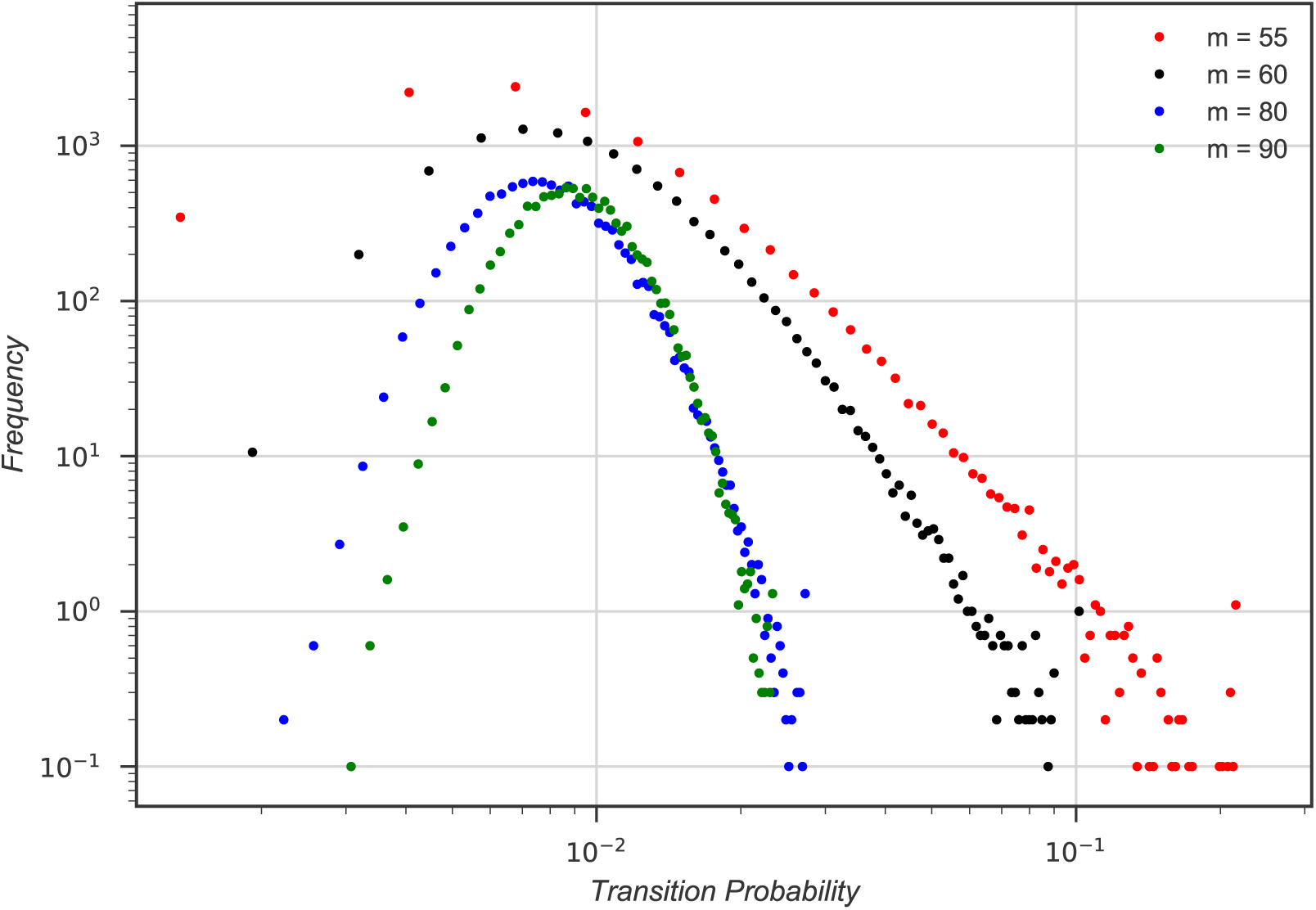
The histogram of transition probability for different densities in the disordered phase in the network of 101 nodes. The histogram peak tends to 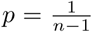 by increasing *x*. It gets close to the power law function as the walker density leans towards *x* = 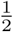.

### transition phase

For *x* = 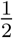, the system would not gravitate towards either of the aforementioned attracting phases. In other words, neither the strengthening nor the weakening role prevails, and as in fact half the containers are empty and the other half occupied, the occupation and availability probabilities of a selected node are equal. In fact, the phase transition takes place exactly when there is an odd number of nodes and the number of the walkers is as follows:

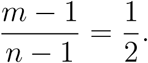

In Fig.5, the transition probability histogram is shown for a network with *n* = 501 and *m* = 251. As it can be seen, the transition probability histogram is scale-free, which indicates that for *x* = 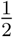, the system will face a critical point. due to the fact that for *x* = 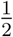 an equilibrium is established between *LTD* and *LTP*, which leads the system to its critical point.

**Figure 5.**
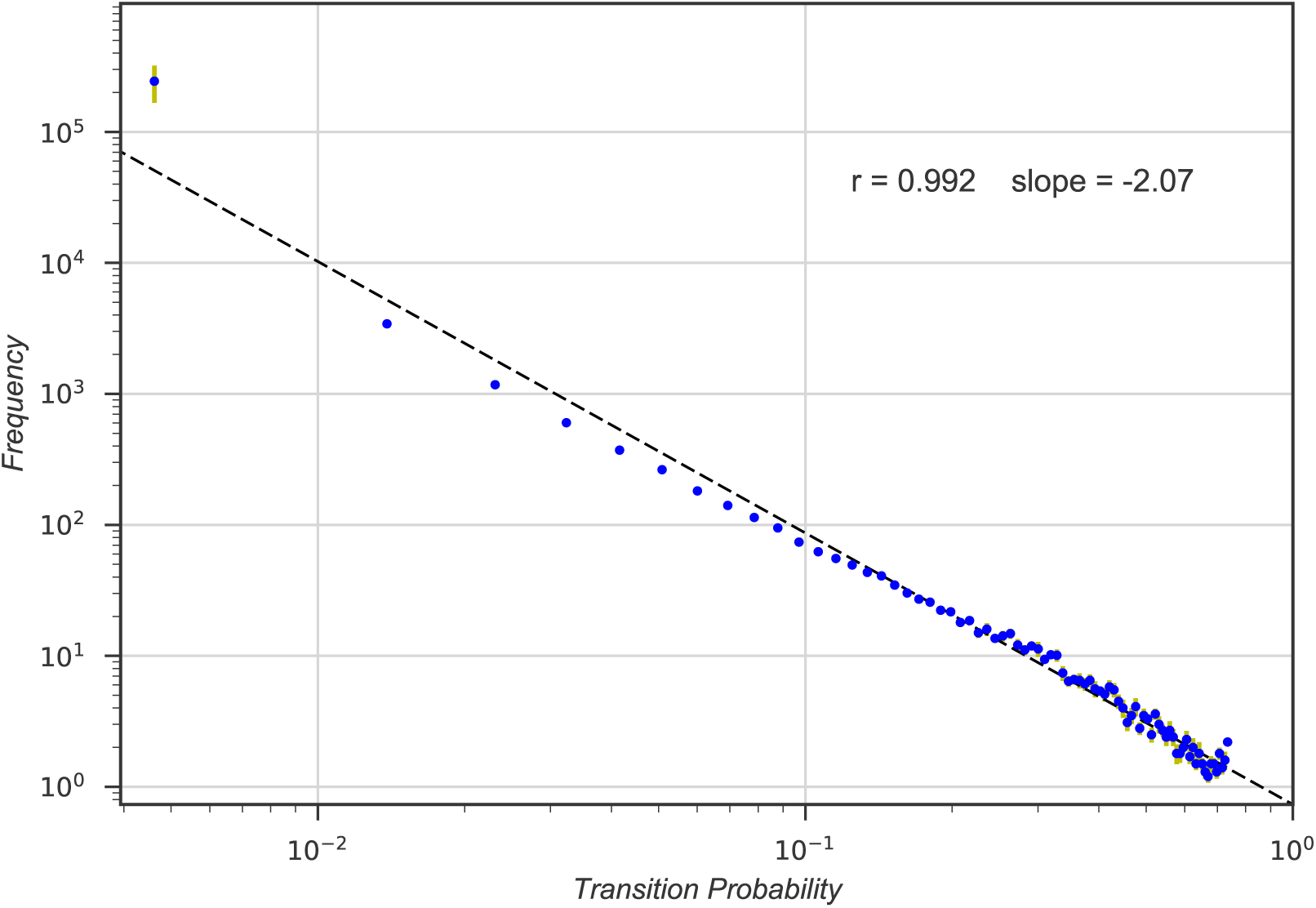
The histogram of weights (transition probabilities) for a network of 501 nodes and 251 walkers. A power law histogram shows that the system is scale free at its transition phase. The fitted power law function (dashed line) has a slope of 2:14 with *r* = 0:992

As previously mentioned, the final state of the system is entirely independent of the initial conditions. In order to put that to the test, we changed the number of the walkers after the system reached its steady state, the system immediately changes its phase to one that is related to its new number of walkers (Fig.6).

**Figure 6.**
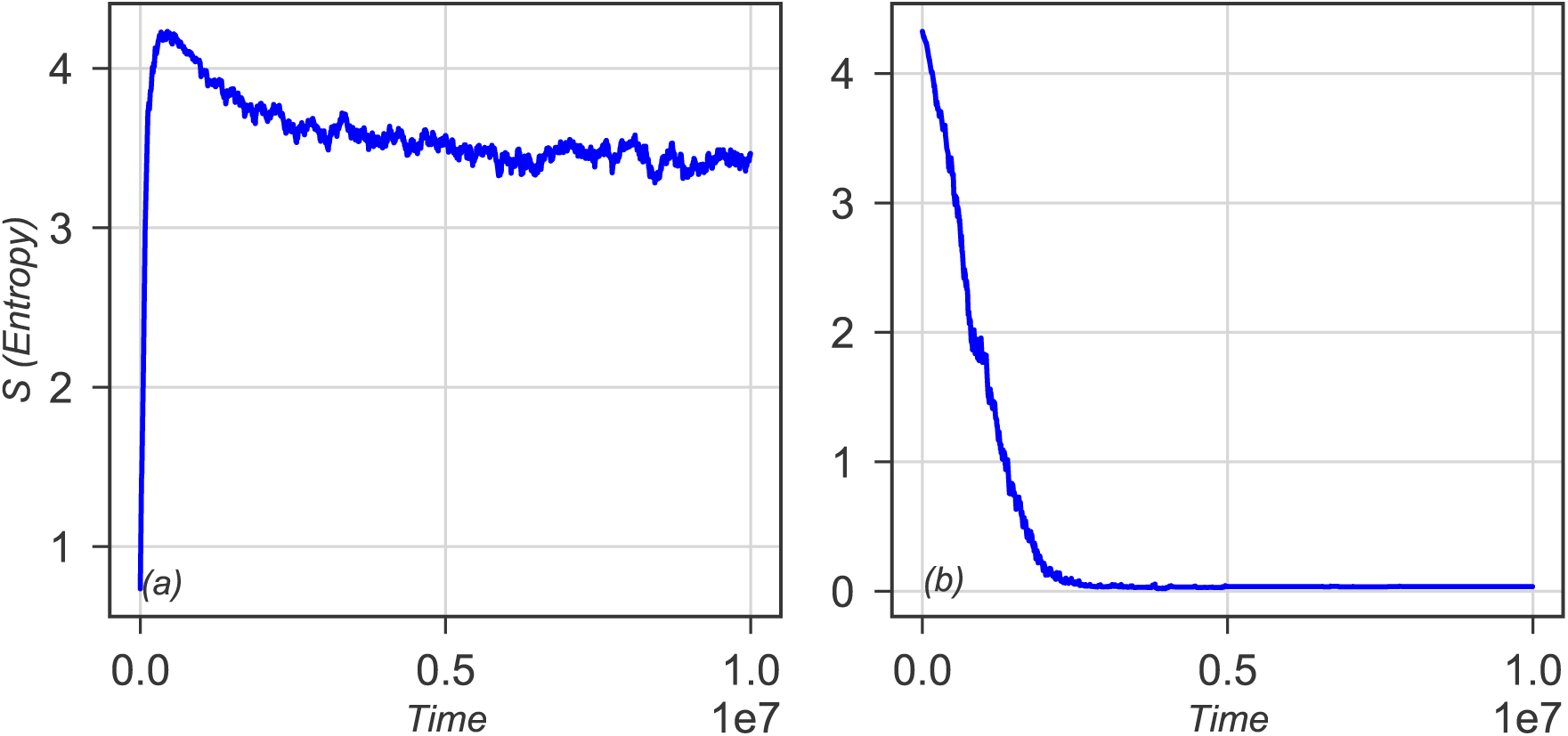
The evolution of the entropy of the system with the initial conditions of a) *x* = 0.51 for a network which has reached the equilibrium with a density of *x* = 0.45. b)*x* = 0.49 for a network which has reached the equilibrium with a density of *x* = 0.55. The entropy of the system rapidly changes so that it corresponds to its current density of walkers. The number of the nodes in both systems are 101.

It is also possible to diagnose the phase of the system from its dynamics, as is shown in Fig.7, for the transition phase, the oscillations of the system past reaching equilibrium are much more frequent compared to the other two phases. Also, there are oscillations in the disordered phase subsequent to the equilibrium that does not emerge in the ordered phase, and this freezing of entropy is a result of the emergence of loops and the walkers trapped inside them.

**Figure 7.**
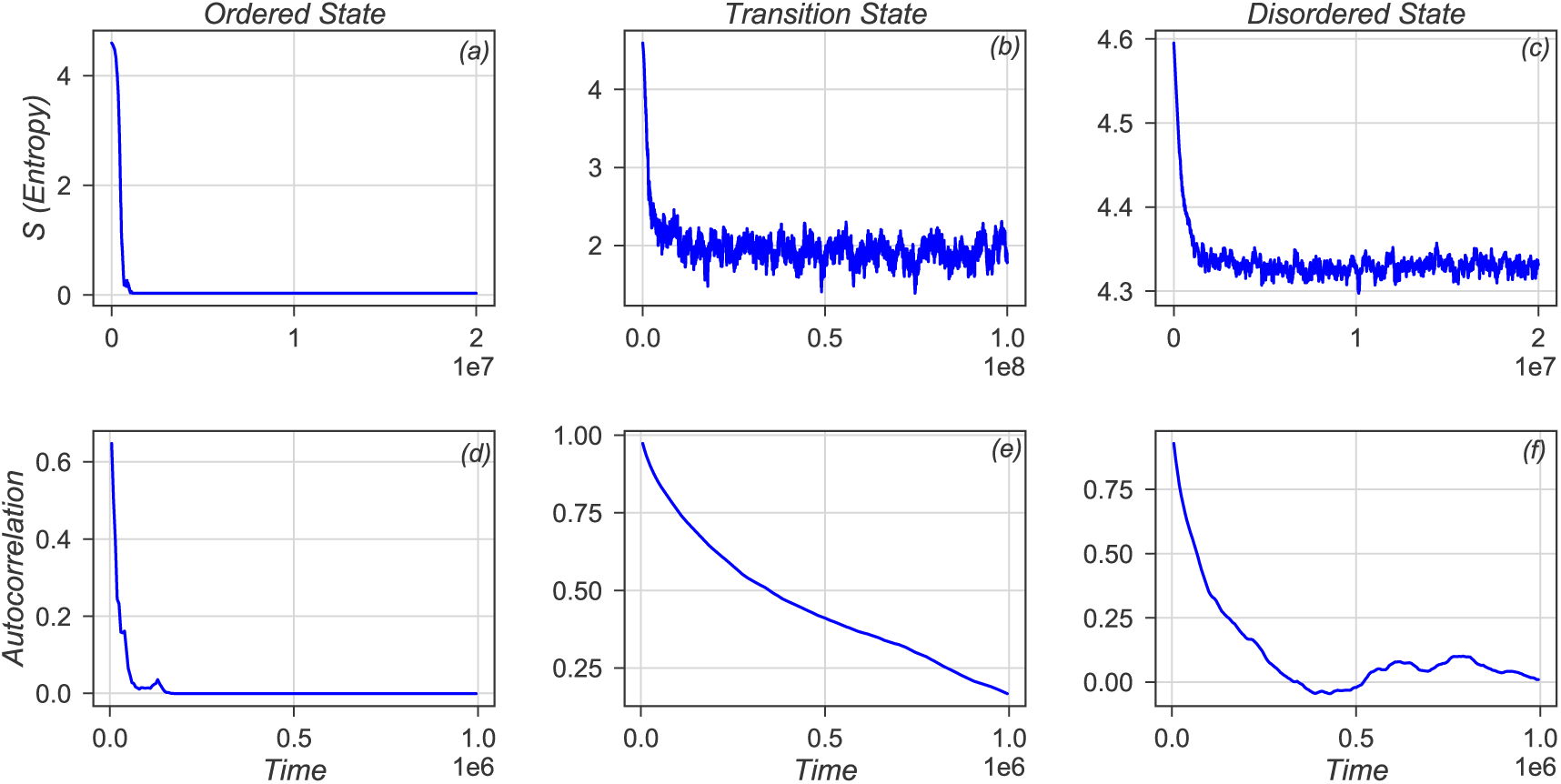
Dynamic of the network entropy and its autocorrelation function for a) and d) ordered phase with 240 walkers, b) and e) transition phase with 251 walkers, c) and f) disordered phase with 450 walkers of a network with 501 nodes. Compared to the other two phases, the system reaches equilibrium much slower in the transition phase, and even after 3×10^8^ time-steps, it has significant fluctuations. Autocorrelation function calculated when the mean entropy of the system remains constant.

To further investigate the steady state of the system as a function of walker’s density, we measured mean clustering coefficient of the network, standard deviation of input strength of nodes, and the mean standard deviation of input and output weights.

Various definitions are available for the clustering coefficient [33, 34, 35]; we used the Zhang-Horvath’s definition [36] that is given with respect to a weighted directed graph(W):

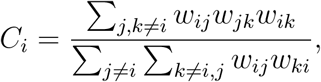

which all weights are normalized by maximum element of *W*. As we expected, when *x* < 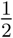 clustering coefficient is small, due to loop formation, and it increases as *x* goes above 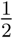, which means all weights tend to be equal (Fig. 8-a).

**Figure 8.**
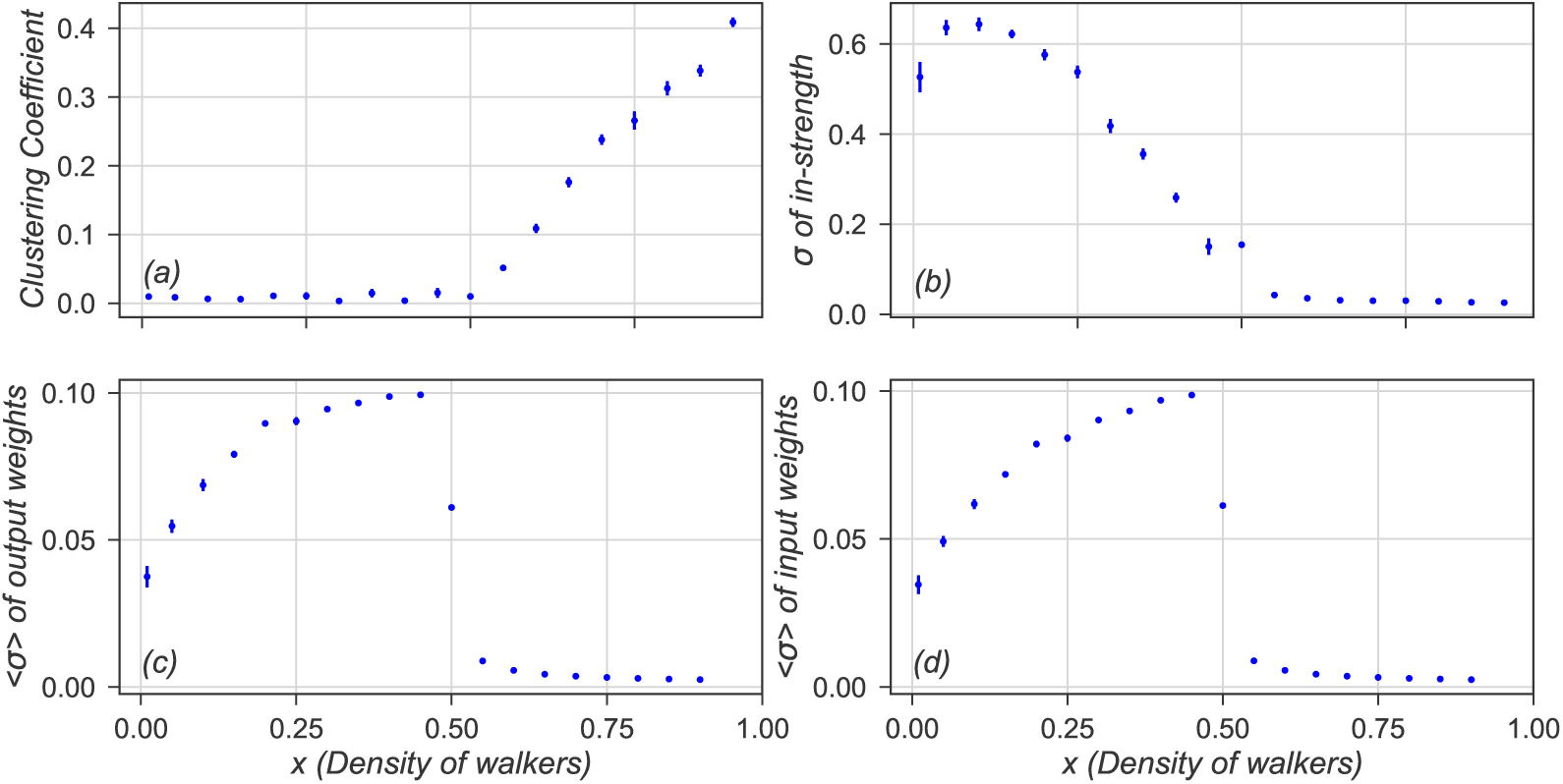
a) Mean clustering coefficient. b) Standard deviation of input strength of nodes. For *x* < 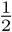, upon decreasing *x* the standard deviation increases, because a few loops form, all walkers will be trapped in them, and that in turn prevents evolving of other links. c) The mean value of the standard deviation of the output weights. d) The mean value of the standard deviation of the input weights, prior to *x* < 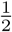,
the standard deviation of both output and input weights is rather large due to loops being formed. For a system with 101 nodes whose every node participates in a loop, this value is equal 0.1, for *x* < 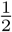, an increase in *x* leads to this value approaching 0.1. Also, for *x* > 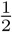, the standard deviation approaches zero, which indicates that all the incoming and also outgoing links are equal. The network size was fixed to 101 nodes, The horizontal axis shows the density of walkers

In-strength and Out-strength of a node are defined by:

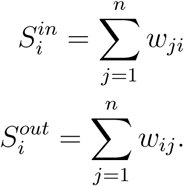

As a result of normalization, the Out-strength for all nodes is equal to 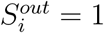, whereas the In-strength is free to take various values. As walkers are neither to be destroyed nor to be created, the average of In-strength over the whole network should be 1. But for different *x*, the standard deviation of the In-strength indicates the similarity among nodes (Fig. 8-b).

Furthermore, the standard deviation of the input (output) weights of a specific node is an indicator of how similar the outgoing (incoming) links to that node are (Fig. 8-c,d).

## 4. Conclusion

In this paper, STDP is modeled using a network evolving with respect to the movement of its reinforced random walkers. We have shown that in the presented model, the steady state of the system is self-organized and it does not depend on the initial conditions of the system (Fig.6), and upon changing the density of walkers, it can fall into three distinct phases: ordered, disordered or transition (Fig.1). We have also shown that the system has a critical point (transition phase), in which the system behaves in a scale free manner, the histogram of transition probabilities (weight of links) is power law(Fig.5) and the relaxation time was too long (Fig.7). As mentioned, the presented model is a simplification of STDP and a lot can be done to improve it. For instance, in a neural network, the activity of the network rises and falls, depending on adaptation, external stimuli or the internal state of the system, that means the number of walkers are not conserved.

## Acknowledgement

We thank to Mohammad Reza Ejtehadi for his fruitful discussions and his critical assessment.

HS thank to Mir Mahmoud Seyed-allaei for the financial support.

## A. Analytical evaluation

According to the Eq. 1 and Eq. 2 we can calculate the conditional expectation value of transition probability for a certain link *ij*, considering *i* is our selected node at time *t* and it already contains a walker:

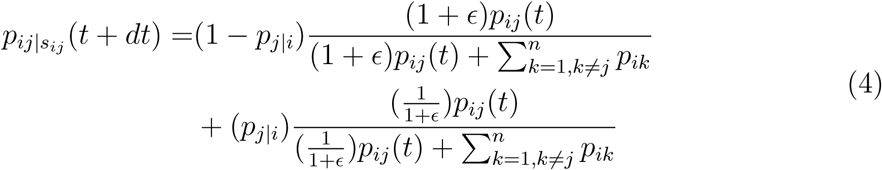

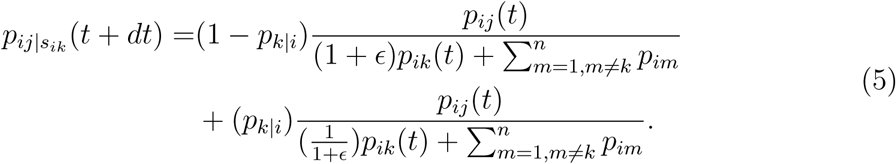

Where *p_ij|s_ik__* (*t* + *dt*) is the conditional expectation value of *p_ij_* considering a walker selected the *ik* link at time *t* and *p_j|i_* is the the probability of existence a walker in node *j* when there is a walker in node *i*. *p_ij_* evolves according to the following equation:

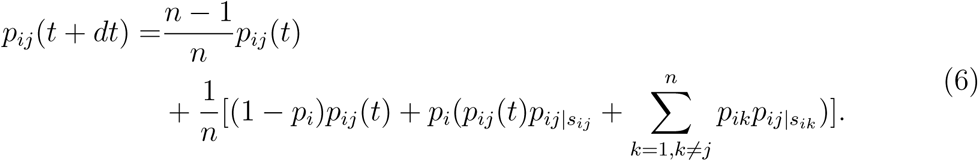

Where *p_i_* is the probability of existence a walker in node *i*. It should be noted that the probability of selecting a certain node is constant and equal to 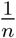. Due to the Eq.4, 5, 6, and with considering the first non-zero terms of *ϵ* we will have:

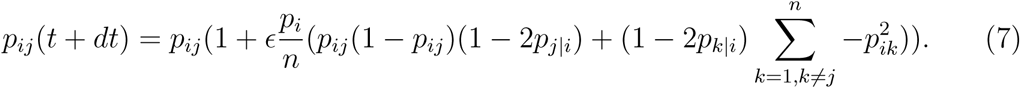

After a sufficiently long time (*t* → ∞), the probability existence of walker remains constant and is equal for all the nodes. Then:

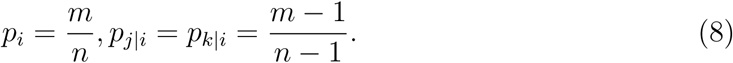

Using Eq.7, 8 transition probability evolves according to:

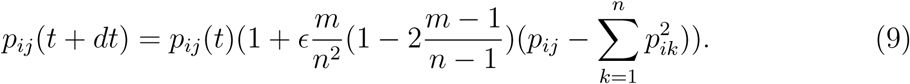

In the stationary state *p_ij_* (*t* + *dt*) = *p_ij_* (*t*), hence the second part of the right hand side of Eq.9 becomes zero:

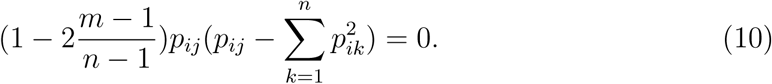

Eq.10 has two equilibrium points :

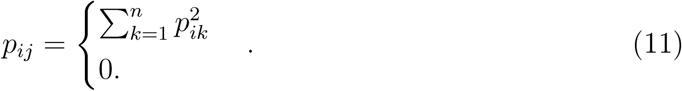

Also we know that :

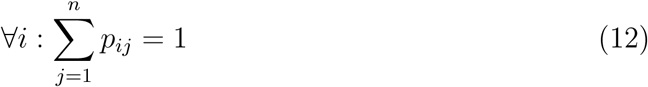

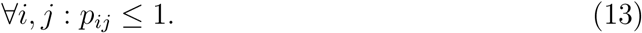

Due to the Eq.11, 12, and 13 our system has two stationary states [21]:

i. Disordered sate: non-zero answer of Eq. 11 is satisfied for all destination nodes which are connected to node 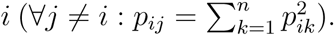 According to normalization (Eq.12) we have:

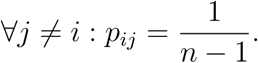
ii. Ordered state: there exists a node k which satisfies non-zero answer of Eq. 11 and for all other destination node, zero-answer of Eq.11 holds. Considering Eq.12 we have:

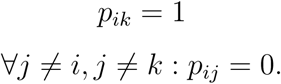

Second order derivative of *p_ij_* determines the stability of the stationary states, due to the Eq.9:

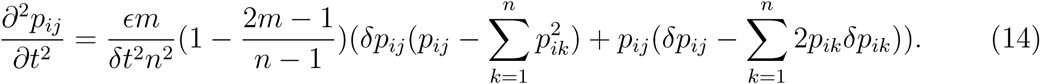

If 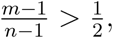 Eq.14 will be greater than zero in the ordered state (so this fixed point is unstable in this regime), and less than zero in the disordered state (so this fixed point is stable in this regime). Hence, for 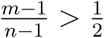, our network reaches the disordered state.

If 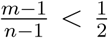 Eq.14 will be less than zero in ordered state and greater than zero in disordered state. Hence, for 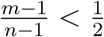, our network reaches the ordered state.

Finally, for 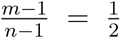, Eq.10 is zero for all conditions and with respect to the simulation results, at this point, the network operates on a critical point.

## References

[1] Timothy V Bliss and Graham L Collingridge. A synaptic model of memory: long-term potentiation in the hippocampus. Nature, 361(6407):31, 1993.

[2] SF Cooke and TVP Bliss. Plasticity in the human central nervous system. Brain, 129(7):1659–1673, 2006.

[3] Wulfram Gerstner, Werner M Kistler, Richard Naud, and Liam Paninski. Neuronal dynamics: From single neurons to networks and models of cognition. Cambridge University Press, 2014.

[4] Yang Dan and Mu-Ming Poo. Spike timing-dependent plasticity: from synapse to perception. Physiological reviews, 86(3):1033–1048, 2006.

[5] Natalia Caporale and Yang Dan. Spike timing–dependent plasticity: a hebbian learning rule. Annu. Rev. Neurosci., 31:25–46, 2008.

[6] D Hebb. The organization of behavior john wiley & sons. New York, 1949.

[7] Henry Markram, Joachim Lübke, Michael Frotscher, and Bert Sakmann. Regulation of synaptic efficacy by coincidence of postsynaptic aps and epsps. Science, 275(5297):213–215, 1997.

[8] Guo-qiang Bi and Mu-ming Poo. Synaptic modifications in cultured hippocampal neurons: dependence on spike timing, synaptic strength, and postsynaptic cell type. Journal of neuroscience, 18(24):10464–10472, 1998.

[9] Per Jesper Sjöström, Gina G Turrigiano, and Sacha B Nelson. Rate, timing, and cooperativity jointly determine cortical synaptic plasticity. Neuron, 32(6):1149–1164, 2001.

[10] Victor Hernandez-Urbina and J Michael Herrmann. Small-world structure induced by spike-timing-dependent plasticity in networks with critical dynamics. arXiv preprint arXiv:1507.07879, 2015.

[11] Sen Song, Kenneth D Miller, and Larry F Abbott. Competitive hebbian learning through spiketiming-dependent synaptic plasticity. Nature neuroscience, 3(9):919–926, 2000.

[12] Richard Kempter, Wulfram Gerstner, and J Leo Van Hemmen. Intrinsic stabilization of output rates by spike-based hebbian learning. Neural computation, 13(12):2709–2741, 2001.

[13] Baktash Babadi and Larry F Abbott. Pairwise analysis can account for network structures arising from spike-timing dependent plasticity. PLoS Comput Biol, 9(2):e1002906, 2013.

[14] Mojtaba Madadi Asl, Alireza Valizadeh, and Peter A Tass. Dendritic and axonal propagation delays determine emergent structures of neuronal networks with plastic synapses. Scientific Reports, 7, 2017.

[15] Burgess Davis. Reinforced random walk. Probability Theory and Related Fields, 84(2):203–229, 1990.

[16] Edward A Codling, Michael J Plank, and Simon Benhamou. Random walk models in biology. Journal of the Royal Society Interface, 5(25):813–834, 2008.

[17] Angela Stevens and Hans G Othmer. Aggregation, blowup, and collapse: the abc’s of taxis in reinforced random walks. SIAM Journal on Applied Mathematics, 57(4):1044–1081, 1997.

[18] Howard A Levine, Serdal Pamuk, Brian D Sleeman, and Marit Nilsen-Hamilton. Mathematical modeling of capillary formation and development in tumor angiogenesis: penetration into the stroma. Bulletin of mathematical biology, 63(5):801–863, 2001.

[19] B Sleeman and IP Wallis. Tumour induced angiogenesis as a reinforced random walk: modelling capillary network formation without endothelial cell proliferation. Mathematical and computer modelling, 36(3):339–358, 2002.

[20] MJ Plank and BD Sleeman. A reinforced random walk model of tumour angiogenesis and anti-angiogenic strategies. Mathematical Medicine and Biology, 20(2):135–181, 2003.

[21] S Mehraban and MR Ejtehadi. A self-organized graph evolution model with preferential network random walk. arXiv preprint arXiv:1205.7069, 2012.

[22] John M Beggs and Dietmar Plenz. Neuronal avalanches in neocortical circuits. Journal of neuroscience, 23(35):11167–11177, 2003.

[23] Dante R Chialvo. Emergent complex neural dynamics. Nature physics, 6(10):744–750, 2010.

[24] Viola Priesemann, Michael Wibral, Mario Valderrama, Robert Pröpper, Michel Le Van Quyen, Theo Geisel, Jochen Triesch, Danko Nikolić, and Matthias HJ Munk. Spike avalanches in vivo suggest a driven, slightly subcritical brain state. Frontiers in systems neuroscience, 8, 2014.

[25] Robert Legenstein and Wolfgang Maass. Edge of chaos and prediction of computational performance for neural circuit models. Neural Networks, 20(3):323–334, 2007.

[26] Ariadne de Andrade Costa, Mauro Copelli, and Osame Kinouchi. Can dynamical synapses produce true self-organized criticality? Journal of Statistical Mechanics: Theory and Experiment, 2015(6):P06004, 2015.

[27] Anna Levina, J Michael Herrmann, and Theo Geisel. Dynamical synapses causing self-organized criticality in neural networks. Nature physics, 3(12):857–860, 2007.

[28] Nigel Stepp, Dietmar Plenz, and Narayan Srinivasa. Synaptic plasticity enables adaptive self tuning critical networks. PLoS Comput Biol, 11(1):e1004043, 2015.

[29] John M Beggs and Nicholas Timme. Being critical of criticality in the brain. Frontiers in physiology, 3:163, 2012.

[30] Heinz Georg Schuster. Criticality in neural systems. John Wiley & Sons, 2014.

[31] Jean-Bernard Brissaud. The meanings of entropy. Entropy, 7(1):68–96, 2005.

[32] Larry F Abbott and Sacha B Nelson. Synaptic plasticity: taming the beast. Nature neuroscience, 3:1178–1183, 2000.

[33] Michael P McAssey and Fetsje Bijma. A clustering coefficient for complete weighted networks. Network Science, 3(02):183–195, 2015.

[34] Duncan J Watts and Steven H Strogatz. Collective dynamics of small-worldnetworks. nature, 393(6684):440–442, 1998.

[35] Wen-Xu Wang, Bing-Hong Wang, Bo Hu, Gang Yan, and Qing Ou. General dynamics of topology and traffic on weighted technological networks. Physical review letters, 94(18):188702, 2005.

[36] Bin Zhang, Steve Horvath, et al. A general framework for weighted gene co-expression network analysis. Statistical applications in genetics and molecular biology, 4(1):1128, 2005.

